# Diet-induced obesity mediated through Estrogen-Related Receptor α is independent of intestinal function

**DOI:** 10.1101/2024.07.10.602978

**Authors:** Kiranmayi Vemuri, Jahangir Iqbal, Sneha Kumar, Alexandra Logerfo, Michael P. Verzi

## Abstract

Obesity has become an epidemic, prompting advances in therapies targeting this condition. Estrogen-related receptor α (ESRRA), a transcription factor, plays pivotal roles in energy metabolism across diverse tissues. Studies have demonstrated that loss of *Esrra* leads to fat malabsorption and resistance to diet-induced obesity. However, the reliance of these studies on germline *Esrra* mutants overlooks the tissue-specific implications of ESRRA in diet-induced obesity. Notably, *Esrra* exhibits high expression in the gastrointestinal (GI) tract relative to other tissues. Given the critical role of the GI tract in dietary lipid metabolism, this study employs mouse genetics and genomics approaches to dissect the specific impact of intestinal ESRRA along with investigating its role in diet-induced obesity.

**Data Transparency:** ChIP-seq and RNA-seq data from this publication have been deposited to GEO accession numbers GSE269824 and GSE269825, respectively. Any additional information required to reanalyze the data reported in this paper is available from the corresponding author upon request.

**Grant Support:** This research was funded by grants from the National Institutes of Health (NIH) to M.P.V. (R01DK121915 and R01DK126446). K.V. was supported by an American Heart Association pre-doctoral fellowship (906006). S.K. was supported by a Rutgers DLS Summer Undergraduate Research Fellowship. A.L. was supported by grants from the NIH grant F31DK137596 and the NIH T32 Biotechnology Training Program (GM135141). The content is solely the responsibility of the authors and does not necessarily represent the official views of the National Institutes of Health.

**Disclosures:** The authors declare no competing interests.

---

There is currently an obesity epidemic and a revolution in therapies targeting obesity. The transcription factor Estrogen-related receptor α (ESRRA) has been implicated both as a mediator of mitochondrial energy metabolism in various tissues [1, 2], and as a regulator of satiety [3]. While loss of *Esrra* has been shown to lead to fat malabsorption and confer resistance to high-fat diet (HFD)-induced obesity [4, 5], a significant limitation in these studies is the use of germline *Esrra* mutants. This approach overlooks the tissue-specific aspects of ESRRA’s involvement in diet-induced obesity. *Esrra* is highly expressed in the GI tract relative to many other tissues (Fig. 1A). Given that the intestine plays a central role in dietary lipid metabolism and is a major target of existing weight loss therapies, such as GLP-1 receptor agonists and orlistat [6], we aimed to investigate whether intestinal ESRRA contributes to diet-induced obesity by using tissue-specific mouse genetic mutants and epigenomic approaches to identify ESRRA target genes in the gut.

**Fig. 1.**
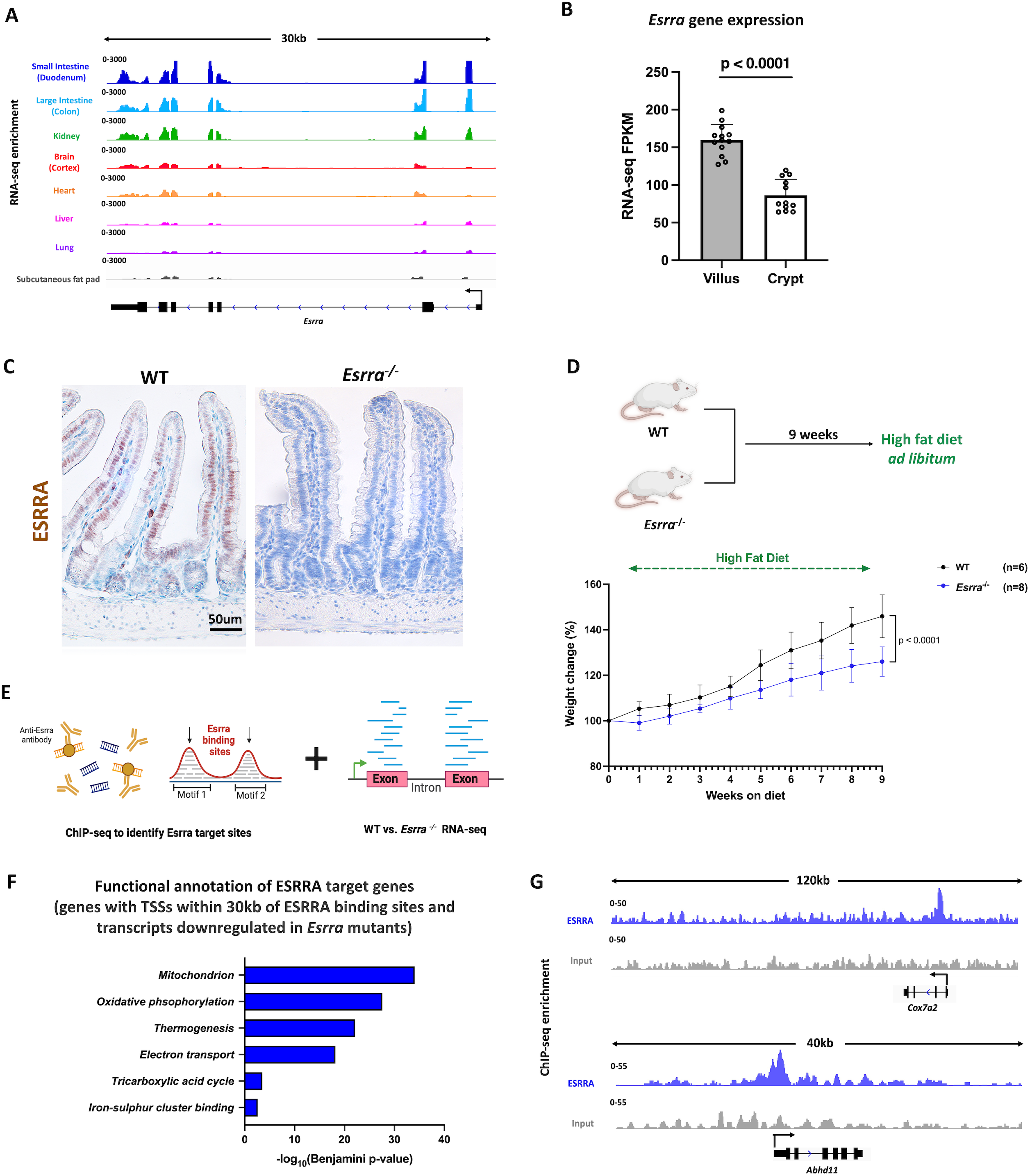
Characterization of intestinal ESRRA function. (A) Transcript expression of *Esrra* across different tissues (GEO: GSE36025). Data visualized using IGV. (B) Bar plots show FPKM values for *Esrra* in crypt and villus derived from RNA-seq (GEO: GSE133949). FPKM data is presented as mean ± s.e.m. (n = 13 villi and 12 crypts, two-sided Student’s *t*-test). (C) IHC staining of ESRRA protein expression in proximal small intestine of WT and *Esrra*^-/-^ mice. Representative of 2 biological replicates per group. (D) Schematic depicting HFD feeding for WT and *Esrra* ^-/-^ mice and body weight changes (%) in *Esrra*^-/-^, and WT littermates upon 9 weeks of HFD feeding (n=6-8 biological replicates per group). (E) Schematic showing integration of ChIP-seq and RNA-seq data to identify ESRRA target genes in the intestinal epithelium. (F) Functional annotation of ESRRA target genes. p-values were calculated using DAVID. (G) Examples of ESRRA binding to gene loci using ESRRA ChIP-seq merged replicates data (RPKM normalized). ESRRA-bound tracks are depicted in blue and input tracks are depicted in gray. Data visualized using IGV.

Within the small intestinal epithelium, we observed that *Esrra* expression was most pronounced in the differentiated villi compared to the proliferating crypts (Figs. 1B, 1C). We hypothesized that ESRRA would be less important in stem/proliferative functions, and enteroids from intestinal crypts of both wild-type (WT) and *Esrra* ^-/-^ mice indeed showed no discernible differences in their growth or budding patterns (Fig. S1A). To validate a role for ESRRA in promoting diet-induced obesity, we fed both WT and *Esrra* ^-/-^ mice a high-fat diet. Consistent with previous reports, we observed a clear resistance to weight gain in the *Esrra* ^-/-^ mice in comparison to their WT littermates, with *Esrra* ^-/-^mice weighing 15-20% less at 9 weeks of feeding (Fig. 1D). We next defined ESRRA target genes in the intestinal epithelium by conducting RNA-seq and ChIP-seq. We compared expression profiles between WT and *Esrra* ^-/-^ intestinal epithelia and identified 526 transcripts downregulated in *Esrra*^-/-^ (as determined by DESeq2 with a log2 fold change < - 0.58 and a p-value < 0.05), which were strongly enriched in annotated mitochondrial functions such as oxidative phosphorylation (Figs. S1B-S1D). This underscores ESRRA’s pivotal role as a major regulator of mitochondrial activity in the intestine, akin to its functions in other tissues [1, 7, 8]. To determine whether ESRRA directly activates these regulated genes, we conducted chromatin immunoprecipitation sequencing (ChIP-seq) and found 1639 genomic regions bound by ESRRA (MACS2 p-value < 10^−5^). ESRRA-bound genomic regions were highly enriched in known ESRR DNA-binding motifs, indicating high-quality and reproducible ChIP (Figs. S1E-S1F). Through integration with RNA-seq data, we identified a set of 110 target genes (defined as bound and regulated by ESRRA as measured by ESRRA ChIP-seq; within 30 kb of ESRRA binding sites; and downregulated in WT vs. *Esrra* ^-/-^ RNA-seq) (Fig.1E). Functional annotation of these direct ESRRA target genes indicated that ESRRA is a major regulator of mitochondrial gene expression in the intestine (Fig. 1F). Examples of two illustrative genes that are direct ESRRA targets and have their products localize to mitochondria are *Abhd11* and *Cox7a2* (Fig. 1G). In summary, these results suggest that ESRRA primarily functions in the differentiated intestinal epithelium as a direct transcriptional activator of genes involved in mitochondrial metabolism.

However, given that the loss of *Esrra* occurs across all tissues (Fig. 1) in this germline knockout model, we couldn’t attribute the observed resistance to diet-induced obesity to intestinal ESRRA. Therefore, we created a tamoxifen-inducible *Esrra* deletion model, integrating *VillinCre*^*ERT2*^ and *Esrra*^*flox/flox*^, to target *Esrra* loss to the intestinal epithelium (Fig. 2A). Intestine-specific loss of *Esrra* (referred to as *Esrra*^*Δ/Δ*^) was confirmed by immunohistochemistry, and tissue pathology was unremarkable (Fig. 2B). We next challenged the *Esrra*^*Δ/Δ*^ mice with HFD alongside littermate *Esrra*^-/-^ and vehicle-treated controls. Interestingly, while we again observed resistance to HFD-induced obesity in *Esrra*^-/-^ mice, the intestine-specific mutants did not exhibit resistance to obesity, suggesting ESRRA in the intestine does not contribute to diet-induced weight gain (Fig. 2C). We next tested whether the mutants exhibited differential feeding behaviors and found that *Esrra*^-/-^ mice consumed less food, while *Esrra*^*Δ/Δ*^ mice consumed similar amounts to controls (Fig. 2D). Consistent with the body weight and feeding behaviors, we noted less white adipose tissue (WAT) in the germline mutant mice (Fig. 2E). Taken together, our findings suggest that intestinal ESRRA does not govern resistance to HFD-induced obesity.

**Fig. 2.**
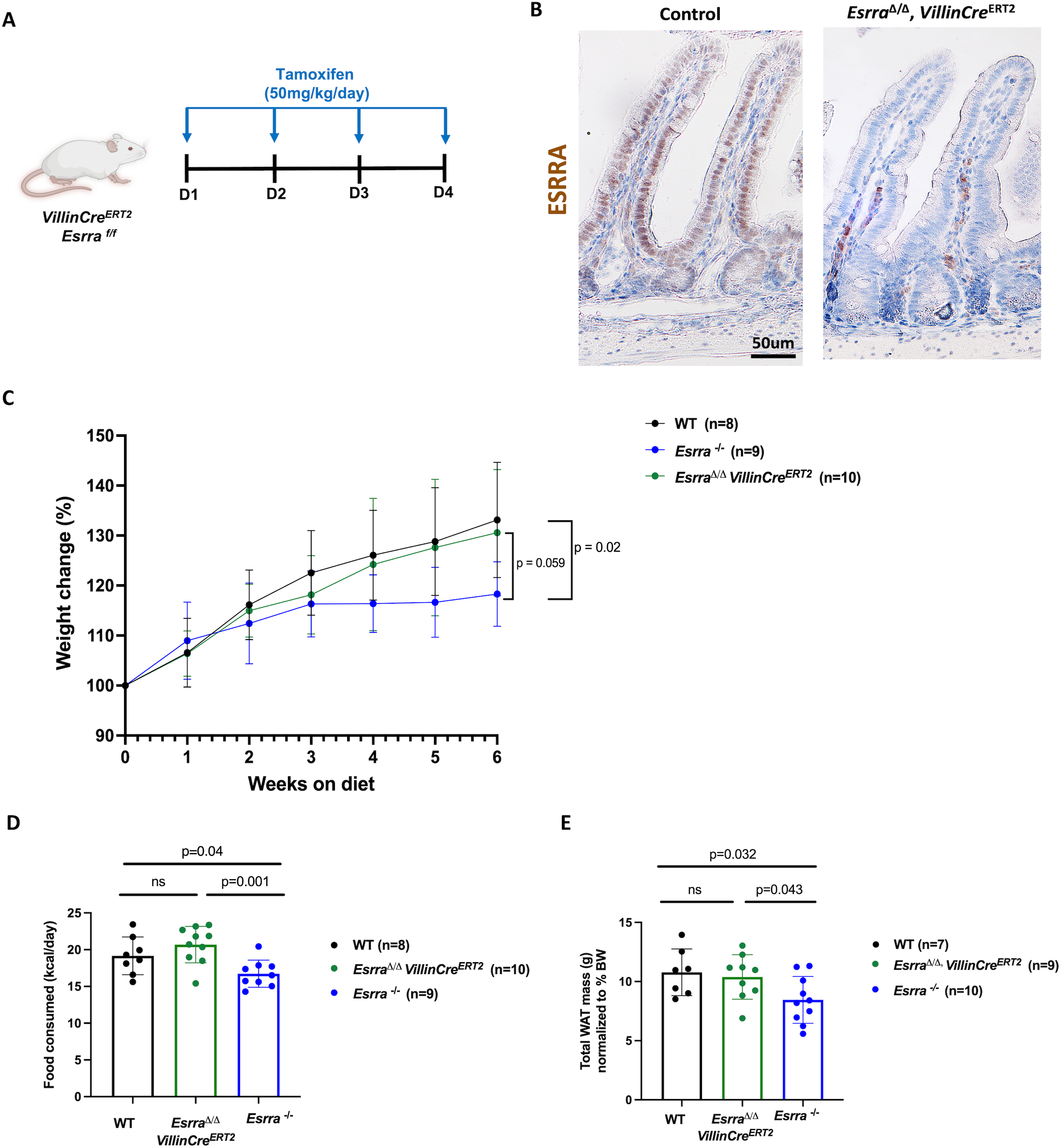
Effect of intestinal ESRRA on resistance to HFD-induced obesity. (A) Schematic illustrating the tamoxifen injection schedule in *Esrra*^*flox/flox*^ mice. (B) IHC staining of ESRRA protein expression in proximal small intestine of WT and *Esrra*^*Δ/Δ*^ mice. Images representative of 2 biological replicates per group. (C) Body weight changes (%) in *Esrra*^-/-^, *Esrra*^*Δ/Δ*^ and control littermate mice upon 6 weeks of HFD feeding. (D) Intake of HFD in *Esrra*^-/-^, *Esrra*^*Δ/Δ*^ and littermate control mice (n=6-8 biological replicates per group). (E) Total white adipose tissue (WAT) mass normalized to body weight in *Esrra* ^-/-^, *Esrra* ^*Δ/Δ*^ and littermate controls (n= 4-6 biological replicates per group). Bars represent mean ± SD.

Our results suggest ESRRA contributes to diet-induced obesity via its role in other tissues such as adipose, muscle, and/or brain. In fact, AAV delivery of shRNA against *Esrra* in the medial pre-frontal cortex leads to reduced feeding and resistance to HFD-induced obesity, suggesting that ESRRA promotes weight gain through behavioral control in the central nervous system [3]. Obesity is a multifactorial condition, influenced by a combination of genetic, environmental, behavioral, and physiological factors. Activation of ESRRA in skeletal muscle can ameliorate diabetes and improve exercise endurance [9], however it might have undesirable effects in various other tissues, as ESRRA is a biomarker for poor prognosis in multiple cancer types [10]. Therefore, it has become increasingly important for future research to delineate the tissue-specific roles of ESRRA and to determine how it can be targeted pharmacologically without causing off-target effects. Our results indicate that therapies targeting ESRRA are unlikely to induce intestine-specific side effects while potentially providing a novel avenue to suppress obesity.

