## Supplementary Information for "Diet-induced obesity mediated through Estrogen-Related Receptor α is independent of intestinal function"

### Materials and Methods:

**Animals:**

*Esrra*$^{f/f}$ (JAX stock #034713) [1] and *Esrra*$^{-/-}$ (MGI:3604552, Jackson Labs) mice were bred to integrate the *Villin-Cre*$^{ERT2}$ transgene [2]. Mice with *Villin-Cre*$^{ERT2}$ and heterozygous for *Esrra*$^{f/-}$ alleles were bred to generate *Esrra*-floxed (*Esrra*$^{f/f}$) and *Esrra*-null (*Esrra*$^{-/-}$) littermate mice. Experimental *Villin-Cre*$^{ERT2}$ *Esrra*$^{f/f}$ mutant mice (6 weeks old) received 4 intraperitoneal injections of tamoxifen at 50 mg/kg/day (Sigma, T5648). Cage mate controls were given corn oil vehicle (Sigma, C8267). All groups were fed high-fat diet (HFD; Research Diets, D12492) *ad libitum*, starting 4 days after the final tamoxifen or oil injection. Mice were weighed weekly in a blinded manner. Following HFD, mice were single-housed for 7 days in wire-bottom metabolic cages to monitor food consumption. Food intake was measured every 48 hours.

For adipose tissue weight measurements, mice were returned to regular cages for 1 week after the metabolic cage period before being sacrificed to collect the duodenum and adipose tissue samples. For RNA-seq experiments, duodenum was collected from *Esrra*$^{-/-}$ and littermate control mice fed HFD for approximately 8 weeks. For ChIP-seq experiments, duodenal villi were collected from C57BL/6 mice fed regular chow. Both male and female mice were used in all *Esrra*$^{-/-}$-related experiments, whereas for experiments involving *Esrra*$^{f/f}$ mutant mice, only males were used. All protocols and experiments were approved by the Rutgers Institutional Animal Care and Use Committee.

**Immunohistochemistry:**

Intestinal tissues were fixed overnight in 4% paraformaldehyde at 4°C, dehydrated through a series of alcohols, xylene, and paraffin embedded. Paraffin sections (5μM thickness) were processed through heat-mediated antigen retrieval in citric acid buffer (pH 6.0) for 30 mins followed by staining with primary antibody (ESRRA, Santa Cruz sc-65718, 1:1,000 dilution). A biotinylated secondary antibody was applied and the Vectastain ABC HRP Kit (Vector Labs, PK-6101) was used for detection. The slides were counterstained with hematoxylin, mounted, and examined using a Lumenera INFINITY3 camera and Infinity Analyze imaging software (v6.5.6).

**Epithelial Cell Isolation:**

The duodenum was dissected and opened longitudinally to wash the lumen and expose epithelial cells. Tissue was cut into 1-inch pieces and rinsed with ice-cold PBS. Tissue was treated with 3mM EDTA/PBS for 35 minutes with the EDTA solution refreshed at the 5- and 10-minute marks. Mechanical force was applied to detach the epithelial cell layer from the underlying mesenchyme. Epithelial cells were either collected in entirety or crypts and villi were collected separately using a 70μM cell strainer. Cells were washed twice with ice-cold PBS and pelleted by centrifugation at 300 rcf at 4°C.

**Intestinal organoid culture:**

Primary crypt-derived organoids from *Esrra*$^{-/-}$ mice and their WT littermate controls were isolated from the duodenum and cultured in Cultrex reduced growth factor matrix R1 (BME-R1)

(Trevigen) according to established methods [3]. Images were captured using a Zeiss Axiovert 200 Microscope (Zeiss, Oberkochen, Germany).

**RNA-seq data analysis:**
Cells were isolated from mouse duodenum as described and processed for RNA extraction using Trizol (Invitrogen, 15596018). RNeasy Micro Kit (Qiagen, 74004) was used to extract RNA according to the manufacturer’s instructions. RNA samples were sent to BGI for transcriptome library preparation and paired-end RNA-sequencing (20 million reads). Raw sequencing reads (fastq) were quality checked using fastQC (v.0.11.3) and were aligned to mouse mm9 reference genome using Kallisto (v2.1.0). DESeq2(v.2.2.1) was used to calculate the fragments per kilobase of transcript per million mapped reads (FPKM) and for differential expression analysis. Genes with FPKM > 1 were used for further analysis. Gene set enrichment analysis (GSEA v.3.0) was performed on the preranked gene list as described previously. Gene ontology analysis was performed with DAVID (v.6.8). RNA-seq data from GEO datasets GSE36025 and GSE133949 was re-analyzed to examine the expression levels of *Esrra* across various tissues

**ChIP-seq and data analysis:**
ChIP-seq was conducted as previously described with slight modifications [4]. Briefly, villus cell pellets were cross-linked in 2 mM DSG (Thermo Scientific, 20593) for 45 min, and then further cross-linked in 1% formaldehyde (Sigma, F8775) for 20 min at room temperature, pelleted and washed with ice-cold PBS. Fixed cells were lysed in 3X volume of lysis buffer (1% SDS, 10mM EDTA, 50mM Tris-HCl pH 8.0, 1X protease inhibitor cocktail (G-Biosciences, 786-433) and sonicated for 8 minutes (30sec on-off, 20% amplitude) using a QSonica Q800R3 sonicator to shear chromatin to 300-600bp. Protein A/G beads (Invitrogen, 10001D and 10004D) were washed with 1% BSA in PBS and pre-loaded with 5μg of anti-Esrra antibody (Santa Cruz, sc-65718). Antibody-conjugated beads were incubated with sheared chromatin in dilution buffer with SDS at a final concentration of 0.24% on a rotator at 4°C overnight. Antibody-conjugated beads were washed five times with RIPA buffer (50mM HEPES pH 7.6, 1mM EDTA, 0.7% Sodium deoxycholate, 1% NP-40, 0.5M LiCl), and once with TE buffer (10mM Tris, 0.1mM EDTA). Cross-links were reversed in 0.1M $NaHCO_3$ and 1% SDS overnight at 65°C. DNA was column purified using the MinElute PCR purification kit (Qiagen, 28004) and quantified using Picogreen (Invitrogen, P7581). Libraries were prepared using the ThruPLEX DNA-Seq kit (Takara, R400675) and DNA Unique Dual Index Kit (Takara, R400666). Paired-end sequencing of ChIP-seq libraries was performed to a depth of 30-50 million reads.

FastQC (v0.11.3) [5], alignment (mm9, bowtie2 (v2.2.6) [6]) and processing were done as previously described [4]. Replicates were merged, sorted and indexed using Samtools (v0.1.19) [7]. deepTools2 bamCoverage (v2.4.2) [8] was used to generate RPKM-normalized bigwig files. The Integrative Genomic Viewer (v2.8.13) [9] was used to visualize normalized bigwig tracks. Enriched ontologies were identified with DAVID (v2021) [10, 11] or GREAT (v4.0.4) [12]. For generation of heatmaps, deepTools2 (v2.4.2) [8] computeMatrix, and plotHeatmap were used.

**Data accessibility:**
ChIP-seq and RNA-seq data from this publication have been deposited to GEO accession numbers GSE269824 and GSE269825, respectively.

**Statistical Analysis:**
Data are presented as mean ± SD or mean ± s.e.m. and graphed using GraphPad Prism (v9.50), with individual replicates plotted. Statistical analyses were conducted with *t*-tests or two-way ANOVA with post hoc analyses, unless otherwise specified.


### Supplementary Figure Legends:

**Fig. S1. Further characterization of intestinal ESRRA function using WT and *Esrra*$^{-/-}$ mice.** (A) Primary organoid cultures isolated from WT and *Esrra*$^{-/-}$ mice. Images are representative of 3 biological replicates. Scale bars are at 500 µm and 50 µm for inset images. (B) Volcano plot of differential gene expression between WT and *Esrra*$^{-/-}$ intestinal epithelial cells (n = 3 biological replicates per group). Differential expression was called with DESeq2 (l2FC > 0.58 or < -0.58, p-value < 0.05). Genes enriched in *Esrra*$^{-/-}$ intestinal epithelia are depicted as blue points and genes enriched in WT-enriched epithelia are depicted as black points. (C) Functional annotation (DAVID) of genes enriched in WT- and *Esrra*$^{-/-}$ intestinal epithelia. p-values were calculated using DAVID. (D) Gene set enrichment analysis (GSEA) shows reduction of genes with mitochondrial functions upon *Esrra* loss in the intestinal epithelium (Kolmogorov-Smirnov test, p < 0.001). (E) Heatmap shows the distribution of ESRRA signal across gene loci for 1639 ESRRA bound sites (MACS2 p-value < $10^{-5}$). (F) ESRRA DNA-binding motifs identified using HOMER. p-values were calculated with HOMER.

**Supplementary Fig. 1**

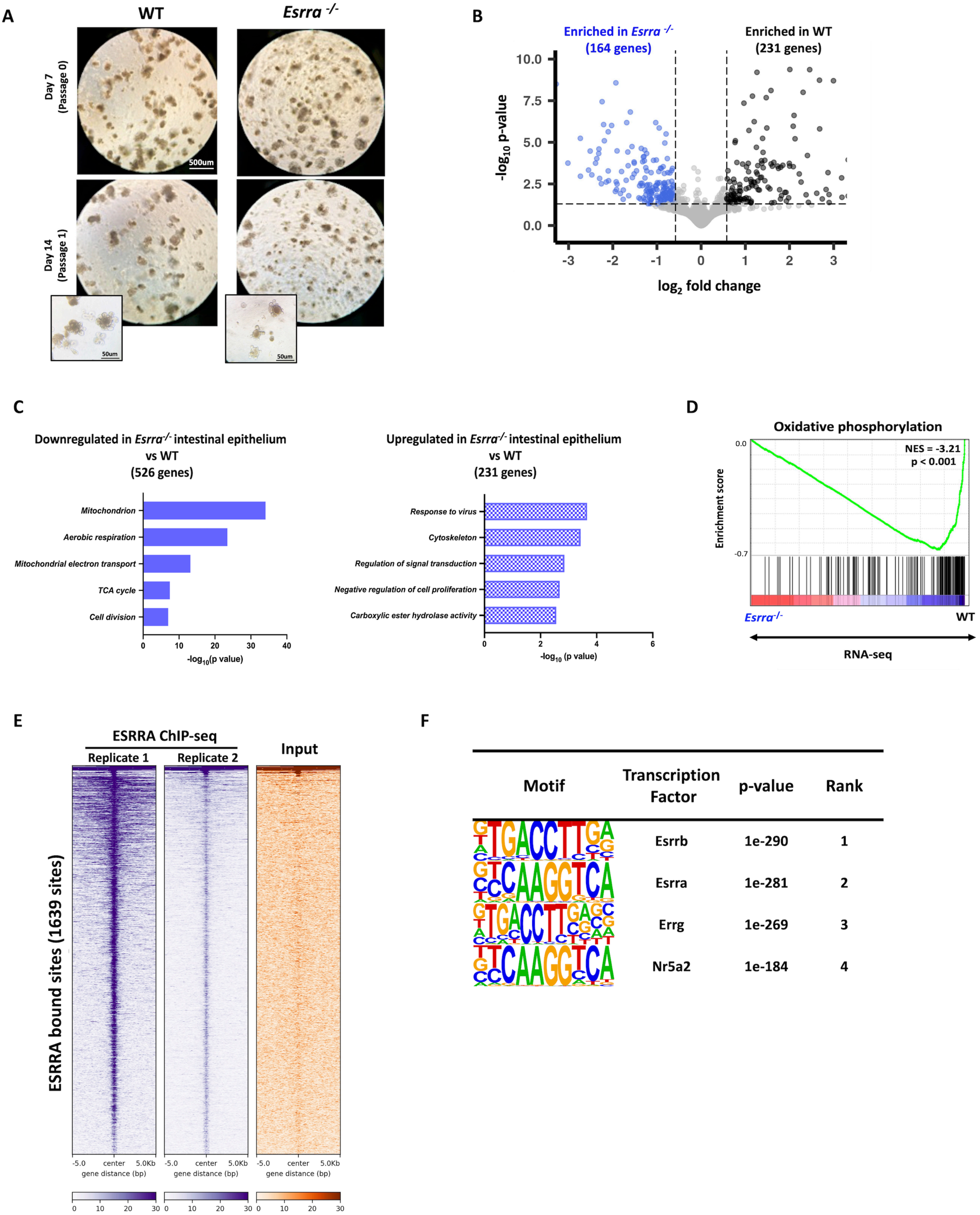
